## Supplementary figures and images for "G protein subunit Gγ13-mediated signaling pathway is critical to the inflammation resolution and functional recovery of severely injured lungs"

### Figure 1- figure supplement 1

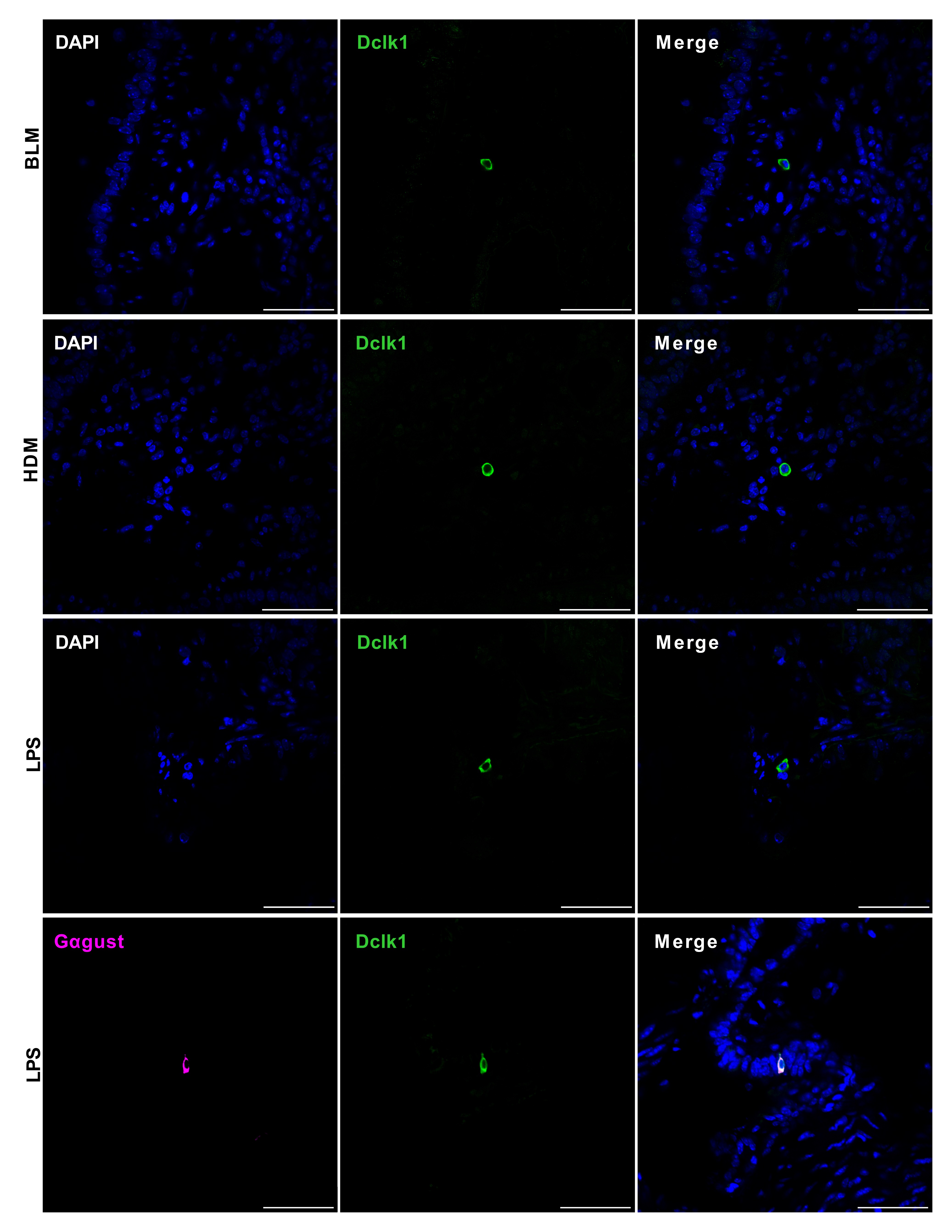

### Figure 2- figure supplement 2

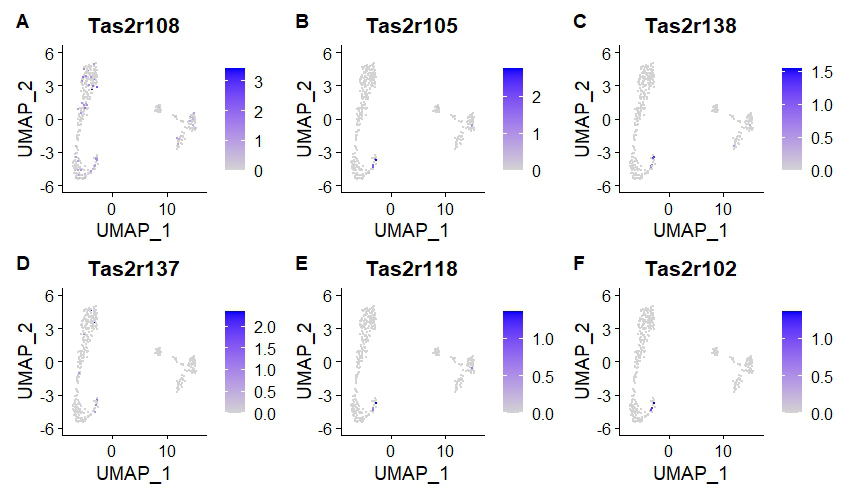

### Figure 2- figure supplement 2

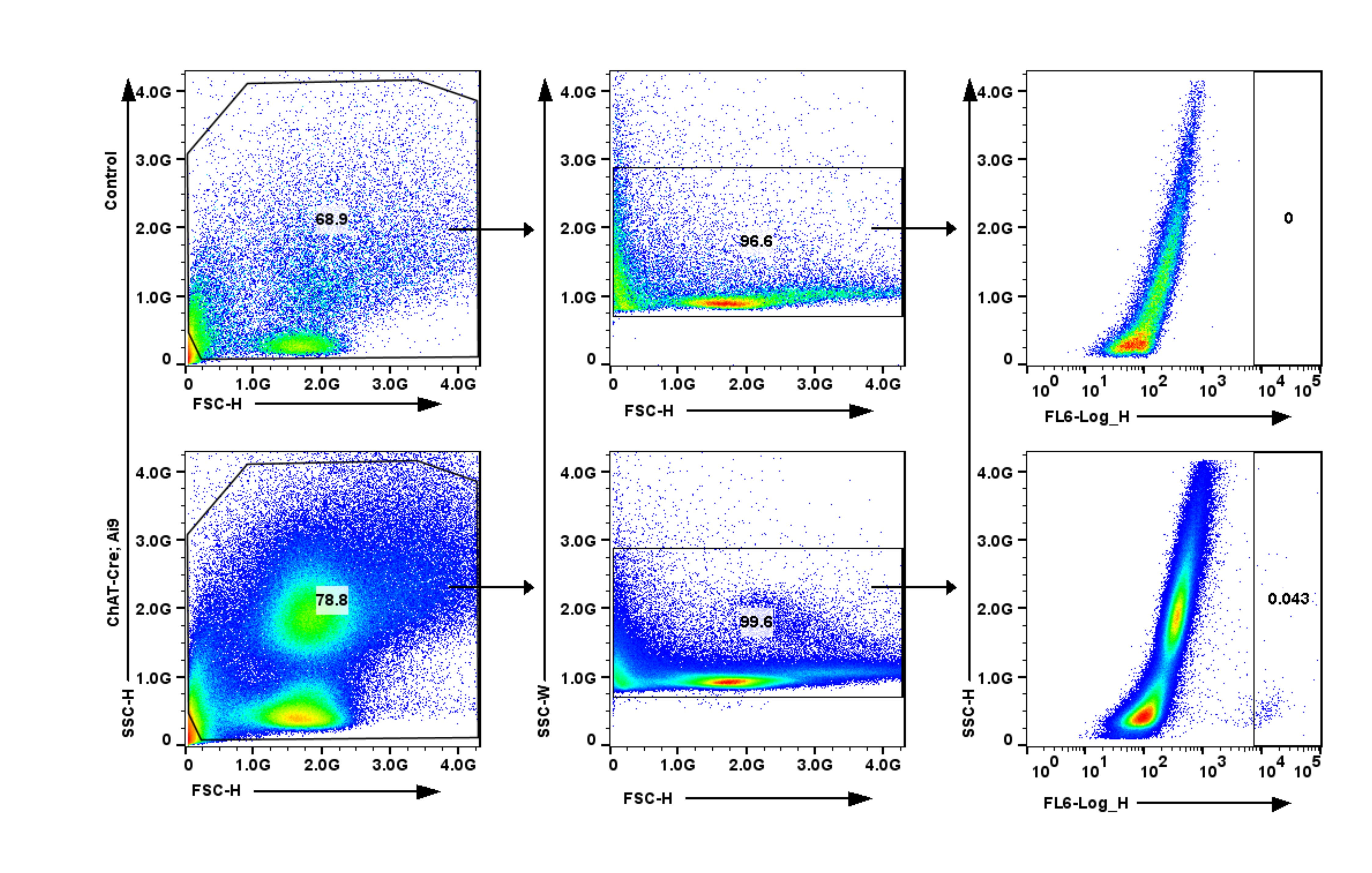

### Figure 3- figure supplement 1

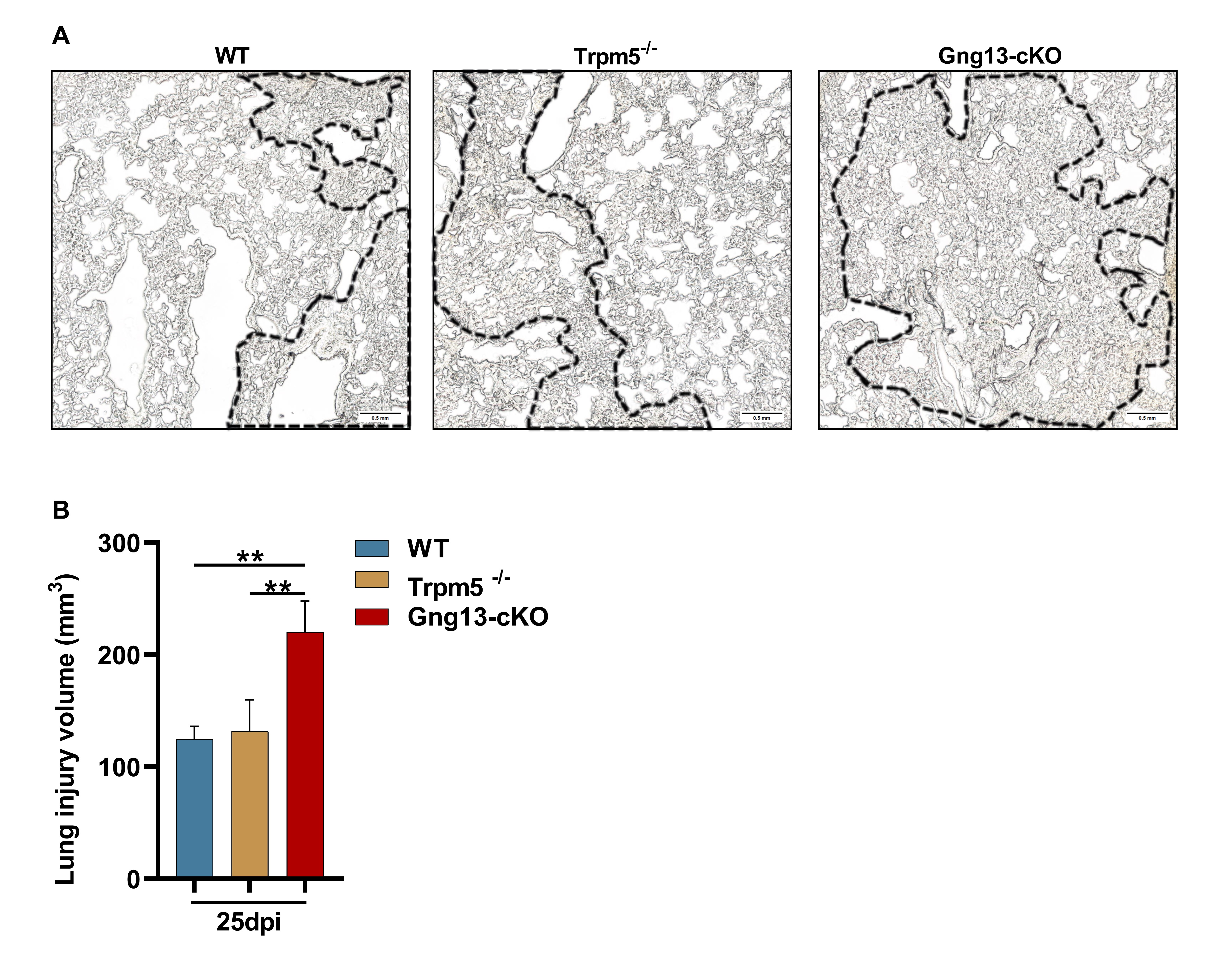

### Figure 3- figure supplement 2

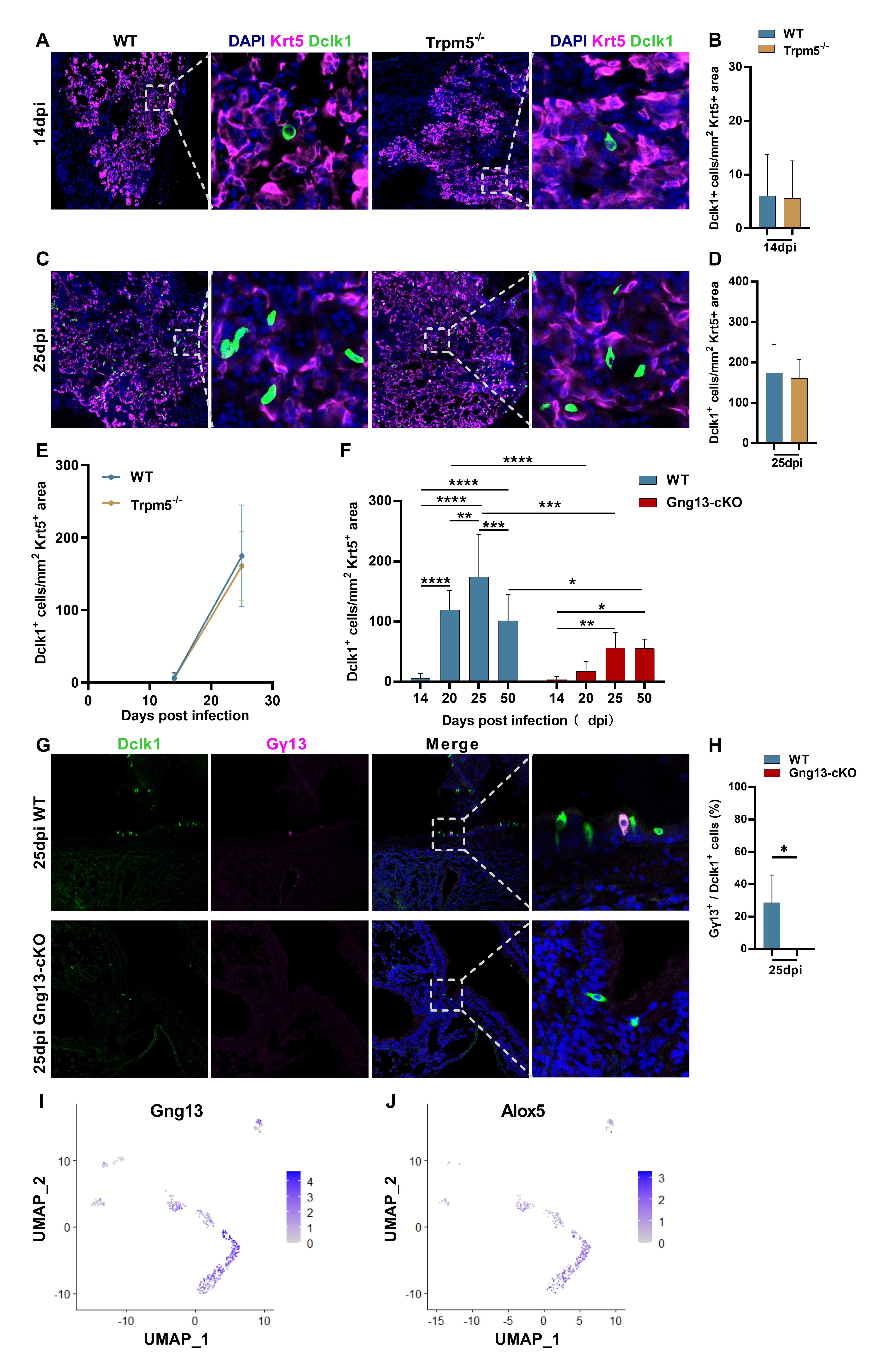

### Figure 4- figure supplement 1

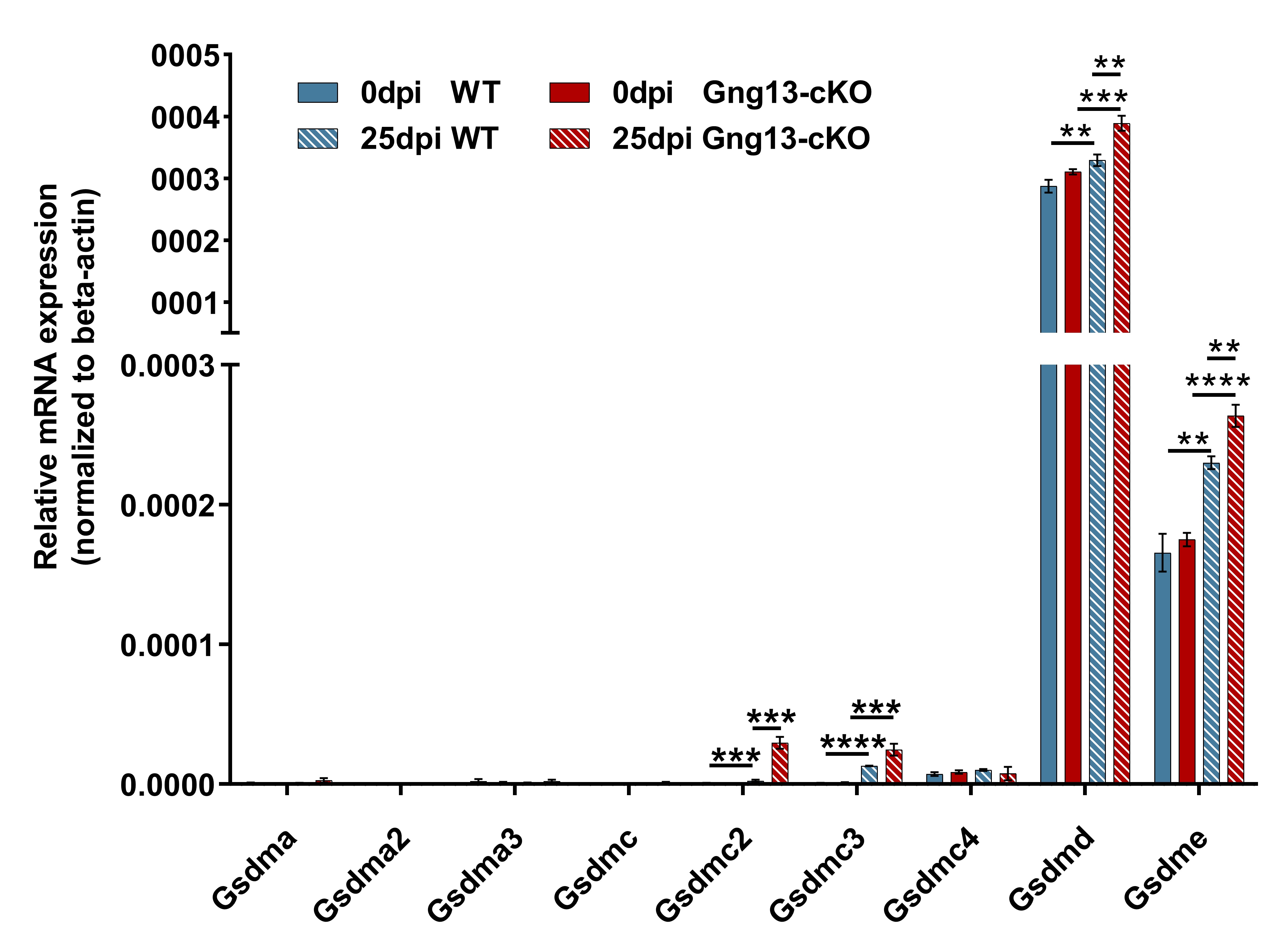

### Figure 5- figure supplement 1

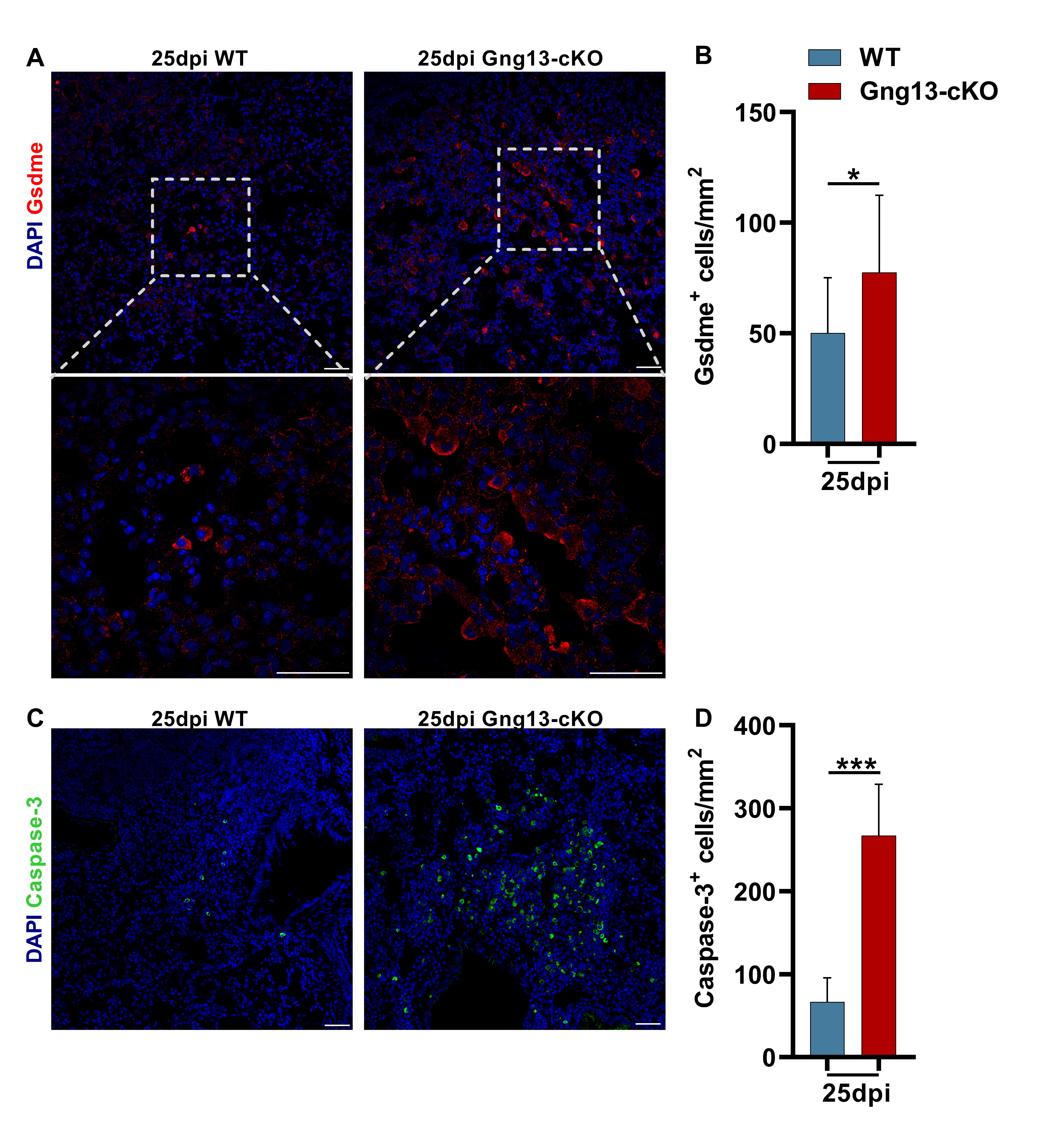

### Figure 5- figure supplement 2

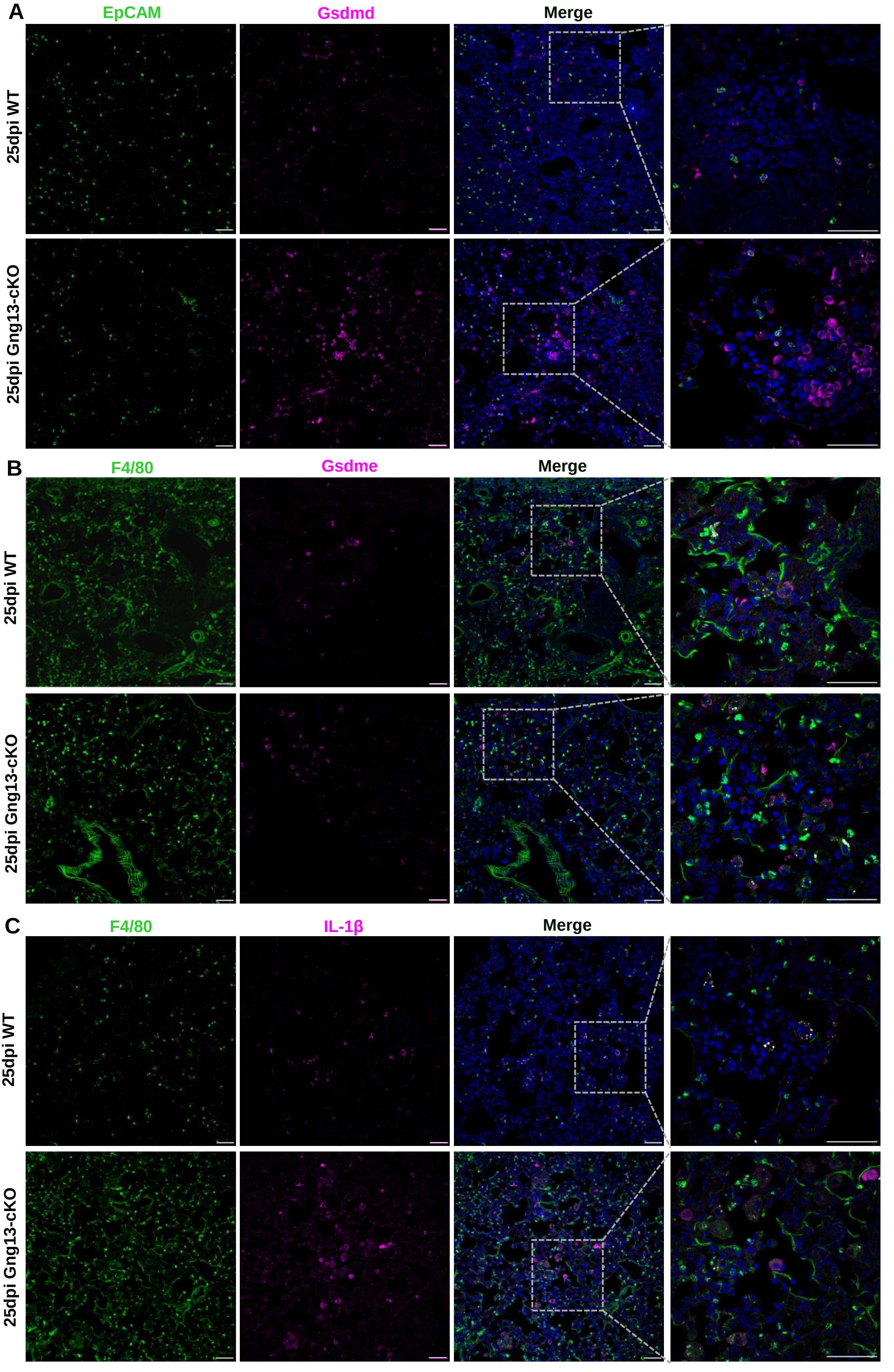
